# THE RHIZOSPHERE BACTERIAL COMMUNITIES DIFFER AMONG DOMESTICATED MAIZE LANDRACES – AN EXPERIMENTAL CONFIRMATION

**DOI:** 10.1101/2021.12.30.474574

**Authors:** Mads Lund, Jacob Agerbo Rasmussen, Jazmín Ramos-Madrigal, M. Thomas P. Gilbert, Christopher James Barnes

## Abstract

- The plant-associated microbiome has been shown to vary considerably between species and across environmental gradients. The effects of genomic variation on the microbiome within single species are less clearly understood, with results often confounded by the larger effects of climatic and edaphic variation.
- In this study, the effect of genomic variation on the rhizosphere bacterial communities of maize was investigated by comparing different genotypes grown within controlled environments. Rhizosphere bacterial communities were profiled by metabarcoding the universal bacterial 16S rRNA v3-v4 region. Initially, plants from the inbred B73 line and the Ancho - More 10 landrace were grown for 12- weeks and compared. The experiment was then repeated with an additional four Mexican landraces (Apachito - Chih 172, Tehua - Chis 204, Serrano - Pueb 180 and Hairnoso de Ocho) that were grown alongside additional B73 and Ancho – More 10 genotypes.
- In both experiments there were significant genotypic differences in the rhizosphere bacteria. Additionally, the bacterial communities were significantly correlated with genomic distance between genotypes, with the more closely related landraces being more similar in rhizosphere bacterial communities.
- Despite limited sampling numbers, here we confirm that genomic variation in maize landraces is associated with differences in the rhizosphere bacterial communities. Further studies that go beyond correlations to identify the mechanisms that determine the genotypic variation of the rhizosphere microbiome are required.

## Introduction

Microbes are associated with nearly all surfaces of plants, and are even present within multiple cell types (1). The rhizosphere is one of the best-studied and most diverse sites of plant-microbe interactions (2). It is the interface between the root and soil, where roots secrete compounds (rhizoexudates) into the adjacent soil altering the abiotic and biotic conditions from the surrounding bulk soil. These secreted rhizoexudates include low molecular weight compounds such as sugars, amino acids, organic acids, phenolics, secondary metabolites, and high molecular weight compounds, such as proteins and mucilage (3,4). They represent a substantial source of carbon and nitrogen input into soil that can be utilised by microbes for growth, and the understanding of the flow of root exudates from plants into microbes (and into the wider soil environment) is vital in understanding carbon, nitrogen, and other nutrient cycles Therefore the quantity and composition of rhizoexudates impacts upon the microbial communities of the rhizosphere, and ultimately the entire ecosystem functioning, with the rhizoexudates from maize specifically found to impact their environment (5). The addition of maize mucilage leads to an increase in nitrous oxide production compared to untreated soils (6), while the addition of artificial maize exudates to soil increased denitrification rates (7). These changes in functioning are derived from changes in microbial composition and biomass, with specific rhizoexudates shown to act as inhibitors, toxins, and repellents of specific microbial species or taxa (5).

There is a growing body of research indicating that the plant genome has a direct and dynamic role in affecting the composition of the microbiome (8,9). For example, a study examining the microbial community composition among genetically diverse maize across five agricultural fields found that the maize genotype significantly influences α- and β-diversity (10). Similar results have been found in experimental plots (9) and greenhouse settings (11), with other plants including of *Arabidopsis thaliana* (the phyllosphere) (12) and barley (between wild and domesticated root communities) (13). A study on the phyllosphere of grapevines also found that the composition of the associated microbiome is correlated with genetic variation of the host (14), suggesting that the aboveground microbial communities can also be influenced by genotypic variation of the host, perhaps also as a result of differing plant exudates from the leaves.

Considering these associations between the genome and microbiome, there are growing calls to consider the host and microbiome together in order to fully understand the host functioning as part of a ‘holobiomic framework’ (15). However, generally variation in the rhizosphere microbiome within species is smaller than between species, and is also considerably smaller than the variation associated with the environment, such as soil pH (16) and average temperature (17). Differences in the plant genotype do not give rise to neat and predictable differences in the microbiome; however there is evidence of more ‘fuzzy’ differences in the composition and functioning associated microbes (analogous to overlapping normal distributions with differing mean averages) (10).

Maize (*Zea mays*, ssp. *mays*) is believed to the result of a long continuous domestication from its wild progenitor teosinte, starting 9,000 years ago in Mexico (18). During this period, traits were selected by people to improve the yield and environmental resilience (19,20). As a consequence of this process, different landraces were produced to match specific but differing local needs (19). However, in the 1970s, the B73 maize inbred line was bred as a result of intense corn breeding programmes (21), representing a particularly high-performing line that led to radically improved corn yields for farmers (22). Its descendants are now present in half the parentage of nearly all hybrid corn around the globe. In this study, we investigated whether the genomic variation between the B73 inbred line and 5 Mexican landraces produced significantly different rhizosphere bacterial communities when grown under common greenhouse conditions (i.e. standardised). This was performed within a glasshouse using the same starting soil to control for environmental variation, where plants were grown for a 12-week period. The rhizosphere bacterial communities were characterised by metabarcoding of the universal 16S rRNA communities. The work was split into two experiments, growth house experiment 1 (GH1) and growth house experiment 2 (GH2). In GH1, the rhizosphere bacteria of the B73 inbred line was compared to a single Mexican landrace (Ancho - More 10). Based on our findings from GH1, an additional experiment was performed that included a further four Mexican landraces (Apachito - Chih 172, Tehua - Chis 204, Serrano - Pueb 180 and Hairnoso de Ocho) (full details of landraces/lines in **Table S1**).

These landraces were selected to have differing geographic ranges and grown in the glasshouse under controlled environmental conditions for a 12-week period. Significant differences in the rhizosphere bacteria were subsequently investigated between the B73 inbred line and Mexican landraces, but also between landraces.

## Materials and Methods

All samples were grown in Frederiksberg growth house (University of Copenhagen, Denmark). Seeds from the B73 inbred line and the Ancho - More 10 [CIMMYTMA 223], Apachito - Chih 172 [CIMMYTMA 7950], Tehua - Chis 204 [CIMMYTMA 794], Serrano - Pueb 180 [CIMMYTMA 1884] and Hairnoso de Ocho [CIMMYTMA 2250] domesticated maize landraces were obtained from the CIMMYT International Maize and Wheat Improvement Center, with the seeds used in GH1 and GH2 were from the same batch (**Table S2**). GH1 occurred between May and July 2019, while GH2 were grown from October to December 2019. For GH1, 3.5L pots were used, which were expanded to 7L volume pots for GH2. Pots were filled with ‘urnejord’ soil, a coarse fertilized and calcium rich soil with a pH of between 5.5 and 6.5. It was composed of: N:140, P:70, K:149, Mg: 247, S: 76, Ca: 2186, Fe:14.4, Mn: 2,8, Cu: 1.7, Zn: 0.9, B: 0.4, Mo: 2.0 g m^-3^. Care was taken to ensure soil was thoroughly homogenised before being added to pots. Seeds were surface sterilised with bleach, before into each pot, four seeds were planted to account for seeds that fail to germinate. Pots were then placed in rows of three to ensure equal light exposure. Temperature was kept to 20 °C, and supplemental light was provided between the hours of 06:00 and 22:00.

Sampling of the rhizosphere was performed after 12 weeks of growth. Plants were pulled from the pot, lightly shaken to remove bulk soil. Roots were then cut from the plant using sterile scissors and placed into DNA free 15 mL tubes and immediately placed at -20 °C. Bulk soil was similarly collected from pots and placed into a DNA free 15 mL tube and immediately placed at -20 °C. At the time of initiating GH2, seeds for each landrace were taken using sterile forceps and placed in 1.5 mL tubes (Eppendorf, Germany) and stored at -20 °C in preparation for DNA extraction.

### Sequencing preparation and sequencing

All DNA extractions were performed in dedicated pre-PCR laboratories within the GLOBE Institute of the University of Copenhagen, Denmark. Between 200-500 mg of roots or soil were extracted from using a soil fast DNA kit (MPBIO). For GH2, single seeds were also extracted (three per genotype), while an extraction negative was included as part of every batch of extractions. Samples initially underwent a mechanical lysis step by adding samples to a Lysing Matrix E tube and placed within a Tissuelyzer II (QIAGEN), which were lysed for 1 minute at 30 Hz. Extractions were then carried out as per manufacturer’s instructions. DNA extracts underwent quantification and quality checking using Qubit Fluorometric Quantification (Thermo Fisher Scientific, Denmark).

As part of library preparation, DNA extracts initially underwent PCR using the 341F (5’-CCTAYGGGRBGCASCAG-3’) and reverse 806R: 5’-GGACTACNNGGGTATCTAAT-3’) primers (23). They target the v3-v4 region of the 16S bacterial gene. They produce an approximate 487 bp amplicon. Each PCR consisted of a 20 µL total reaction volume containing 2 µL of 1x Gold PCR buffer, 2.5mM MgCl_2_, 0.2mM dNTP, 0.1 U Taq Gold polymerase, 0.5 mg mL^-1^ BSA (all of which were from Applied Biosystems, Denmark) and 2 μL of DNA extract. DNA amplification was carried out in a thermal cycler using the following conditions: 95° C for 10 min, followed by 15 s at 95 °C, 20 s at 53 °C, and 40 s at 72 °C. There was a final extension of 7 minutes at 72 °C. In GH1, PCR reactions ran for 23 cycles for the soil samples and 28 cycles for the rhizosphere samples. However, the first PCR amplification of for the rhizosphere extraction in GH1 failed and the PCR amplification was repeated with 33 cycles using the same thermal cycler settings. In the following GH2 experiment, the PCR reaction ran for 23 cycles for the soil samples, 28 cycles for the seed samples and 28 cycles for the rhizosphere samples. All samples from both GH1 and GH2 were ran in triplicate to provide technical replicates, while extraction negatives and PCR negatives (sterile water used as a substitute for DNA extract) were included for all batches of PCR amplifications.

PCR products were pooled into four separate libraries, with 100 ng of DNA suspended in a total volume of 32 µL. Pools then underwent library preparation for sequencing using the Blunt-End Single-Tube (BEST) approach that was specifically modified for metabarcoding (24). Library products were purified with magnetic beads (SPRI beads, Beckman-Coulter, IN, USA) using a 1.2:1 ratio of beads to library volumes to remove unbound adapters. Samples were eluted into 35 µL of EB-buffer (QIAGEN, Germany). Samples were then sequenced with 600 cycles chemistry on the MiSeq v3 sequencing platform (Illumina, CA, USA) at the National High-Throughput Sequencing Centre of Denmark (University of Copenhagen, Denmark).

### Bioinformatic analysis of metabarcoding data

Raw reads were uploaded to the Electronic Research Data Archive (ERDA), hosted by University of Copenhagen under the given DOI https://doi.org/10.17894/ucph.ef1d47d6-7979-474f-b918-cd1897143dc6. Adapters were removed using AdaptorRemoval (v2.3.1) (25) and Cutadapt (v2.3.1) (26) was used to check and correct the orientation of reads. Technical triplicates (three per sample) were manually checked for consistency (using nMDS ordinations) and summed in order to maximise reads per sample. All subsequent data processing were performed using the DADA2 pipeline within the *R* statistical computing environment (27). Reads were merged together with an overlap of 15 bp instead of 20 bp due to long amplicon size. Reads were denoised, chimeras were removed, and taxonomy assigned using the Silva 132 reference database. The post-clustering algorithm LULU (v0.1.0) (28) has been shown to reduce the occurrence of sequence errors and was therefore applied to ASVs to further curate data post ASV inferring.

Potential contaminants were identified and removed using the package decontam (v1.10.9) using a prevalence identification method. Here, the occurrences of ASVs within samples versus negative controls were compared, and a stringent threshold of 0.5 was used as a cut-off for removal.

Reads that were not assigned to the bacterial kingdom were removed. Additionally, reads that are assigned to mitochondria and chloroplast have been shown to wrongly co-amplify with the bacterial primers (29). Numerous reads were found to assign to each, particularly within the seed samples. Sequences were blasted against the maize chloroplast and mitochondria, which were 100% homologous. Therefore, these sequences were also removed. These reads dominated seed samples, and to such an extent, no meaningful bacterial profiles could be made for any seed sample and therefore these samples were disgarded from further analyses. For the remaining samples, rarefaction curves were plotted after filtering, suggesting that an asymptote was almost reached for most samples (**Fig. S1**). Richness was calculated by rarefying to the lowest accepted read depth (1307 reads) and used in downstream analyses. Unrarified read counts underwent a fourth root transformation before being converted into relative abundances for use in analyses of community composition (**Table S3**).

### Genomic distances

To estimate the genomic distance between the maize landraces, we used the sequencing data generated in Swarts et al. 2017 (30). This dataset consists of 1,316 wild and domesticated maize samples that were genotyped using a genotype-by-synthesis strategy and included representatives of the landraces used in this study. We downloaded the raw sequencing reads from the NCBI SRA from relevant accessions: Tarahumara Serrano-ZM22-006 (n=5), Tepehuan Serrano-ZM22-003 (n=3), Mountain Pima Maiz Ancho-ZM11-001 (n=2), Mountain Pima Maiz Ancho-ZM20-003 (n=4), Tarahumara Harinoso de Ocho-ZM11-008 (n=4), Tarahumara Harinoso de Ocho-ZM10-004 (n=4), Apachito Chihuahua-ZM13-017 (n=3), Tarahumara Apachito-ZM13-022 (n=1), Apachito-Mex_High (n=1) and Tehua-Mex_Low (n=1). Adapter sequences, low quality stretches and leading/tailing N’s were trimmed from the raw reads using AdapterRemoval 2.0 (25). Reads 25 bp or shorter after trimming were discarded. Read were mapped to the *Zea mays* ssp. *mays* reference sequence (B73-v3.25) (31) using bwa aln 0.7.12 (32). Reads with a mapping quality below 30, or ambiguously mapping to the reference genome were discarded. Reads mappings were realigned using GATK.3.3 (33) and the MD-tag was recalculated using samtools 1.2 (34).

Given that the average depth of coverage of the samples was ∼1.4x, we sampled a random read per site instead of performing genotype calling. For each sample and each of the SNPs identified in the maize Hapmap2 diversity panel (35) we randomly sampled one allele using ANGSD v0.920 (36) at bases with a minimum quality of 20. Additionally, we incorporated the B73 reference genome to the SNP dataset by taking the base present in the assembly for each of the SNPs. The final dataset consisted of 757,119 SNP sites across the 29 maize accessions. Identity-by-state (IBS) pairwise distances were estimated using plink2.0 (37) (--distace-matrix) on the final dataset.

### Statistical analysis

All analyses were performed using *R* (v5.3.3) and visualised using the ggplot2 package (38), apart from the heat tree, which was created using the Metacoder package (39).

To calculate differences in community composition associated between genotypes and bulk soil samples, PERMANOVAs were performed using the adonis function. Post-hoc analyses were performed using the betadisper function, with Tukey Honest Square Differences being calculated to determine pairwise differences between samples. Community variation was visualised by calculating a Bray-Curtis similarity matrix and performing non-metric multidimensional scaling (nMDS). Differences in bacterial richness between genotype and bulk soil were calculated using Kruskal-Wallis tests (using the richness measures calculated from rarefied data). Post-hoc analyses were performed to calculate pairwise comparisons between samples, which was performed using Dunn tests.

To investigate the relationship between the genomic distances of the landraces to the microbiome, the IBS distance matrix was correlated against the Bray-Curtis similarity matrix for the GH2 rhizosphere samples using a Mantel test and Procrustes ordination.

## Results

### GH1 – The rhizosphere bacteria of the B73 and Ancho - More 10 differed in composition

Initially, significant genotypic variation in the rhizosphere was assessed for between the modern inbred B73 line and a Mexican landrace (Ancho - More 10) within a glasshouse (and therefore without confounding environmental variation). PERMANOVA revealed a significant difference in community composition between the two (*R*^2^ = 0.239, *P* = 0.045), accounting for 23.9% of community variation (**Fig. 2A**). While bacterial richness was higher in the Ancho - More 10 landrace (199.8 ±52.2 ASVs) than the B37 line (163.3 ±22.5 ASVs) (**Fig. 2B**), this was not significant (*H* = 2.08, *P* = 0.149). The B73 line was characterised by a higher abundance of Verrcomicrobia, Actinobacteria and some Gammaproteobacteria (**Fig. 3**), while the Ancho - More 10 landrace had higher abundances of Patescibateria, Dependentiae and Chloroflexi.

**Figure 1.**
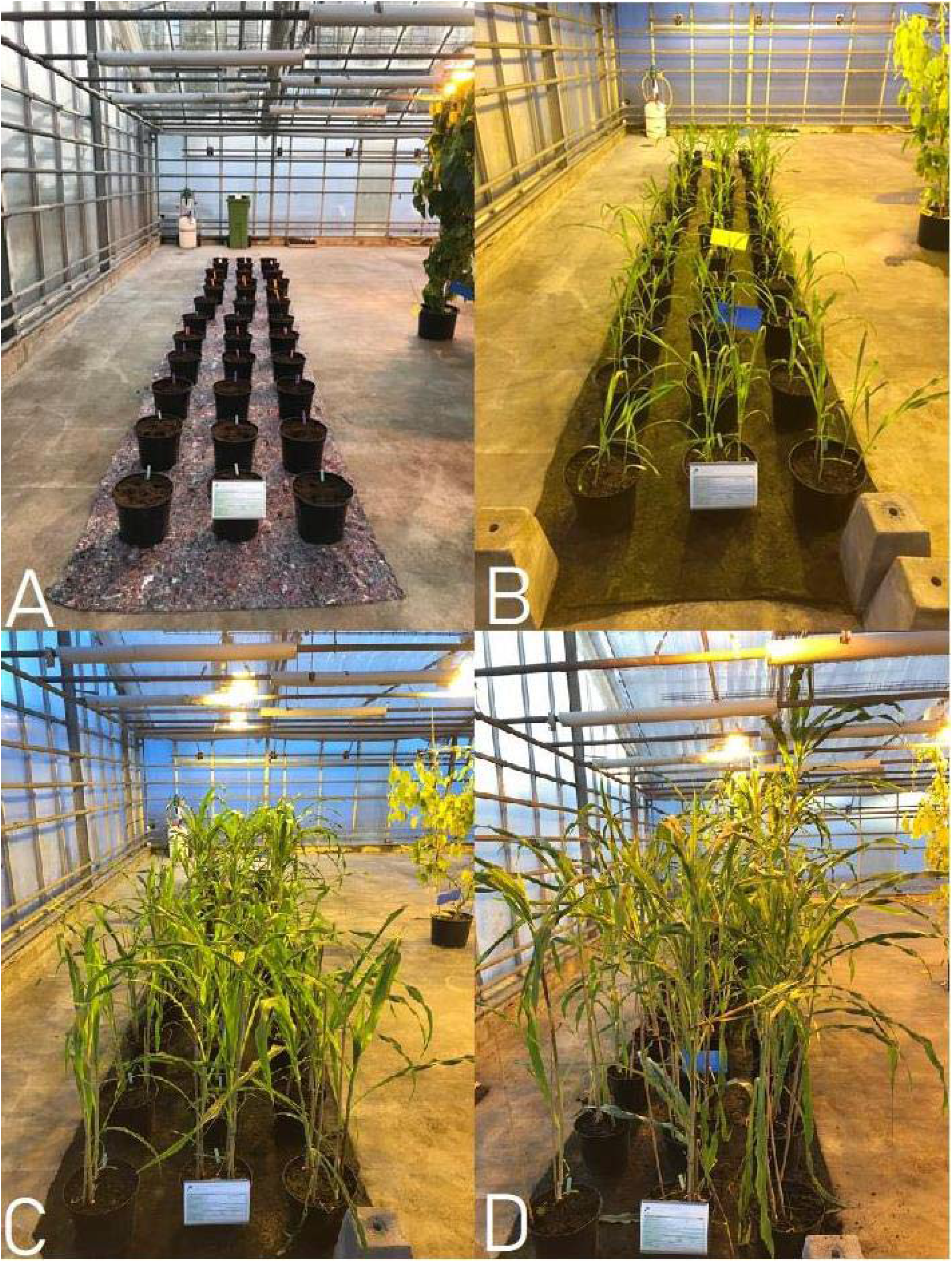
GH1 experimental setup. Maize plants from the B73 inbred line and Ancho - More 10 domesticated landrace were planted (**A**) and measured at 4-weeks (**B**) and 8-weeks (**C**) after planting. Plants were the harvested after 12-weeks (**D**) and their rhizosphere bacteria profiled by DNA metabarcoding.

**Figure 2.**
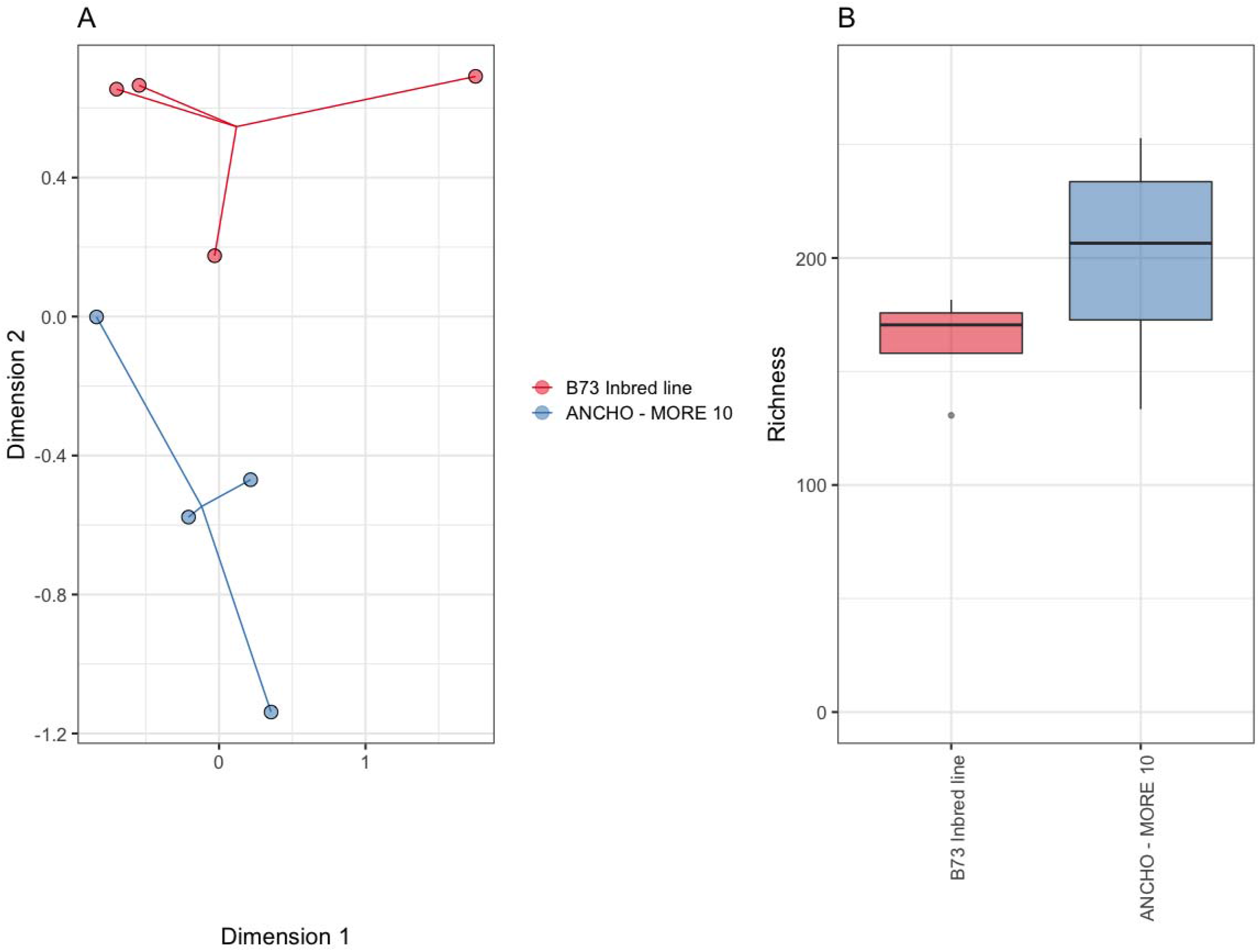
(**A**) Non-metric multidimensional scaling plot visualised significant differences between the rhizosphere bacteria from the inbred B73 line and Ancho - More 10 Mexican landrace from GH1. (**B**) While there were more ASVs on average within the Ancho - More 10 landrace, it was non-significant.

**Figure 3.**
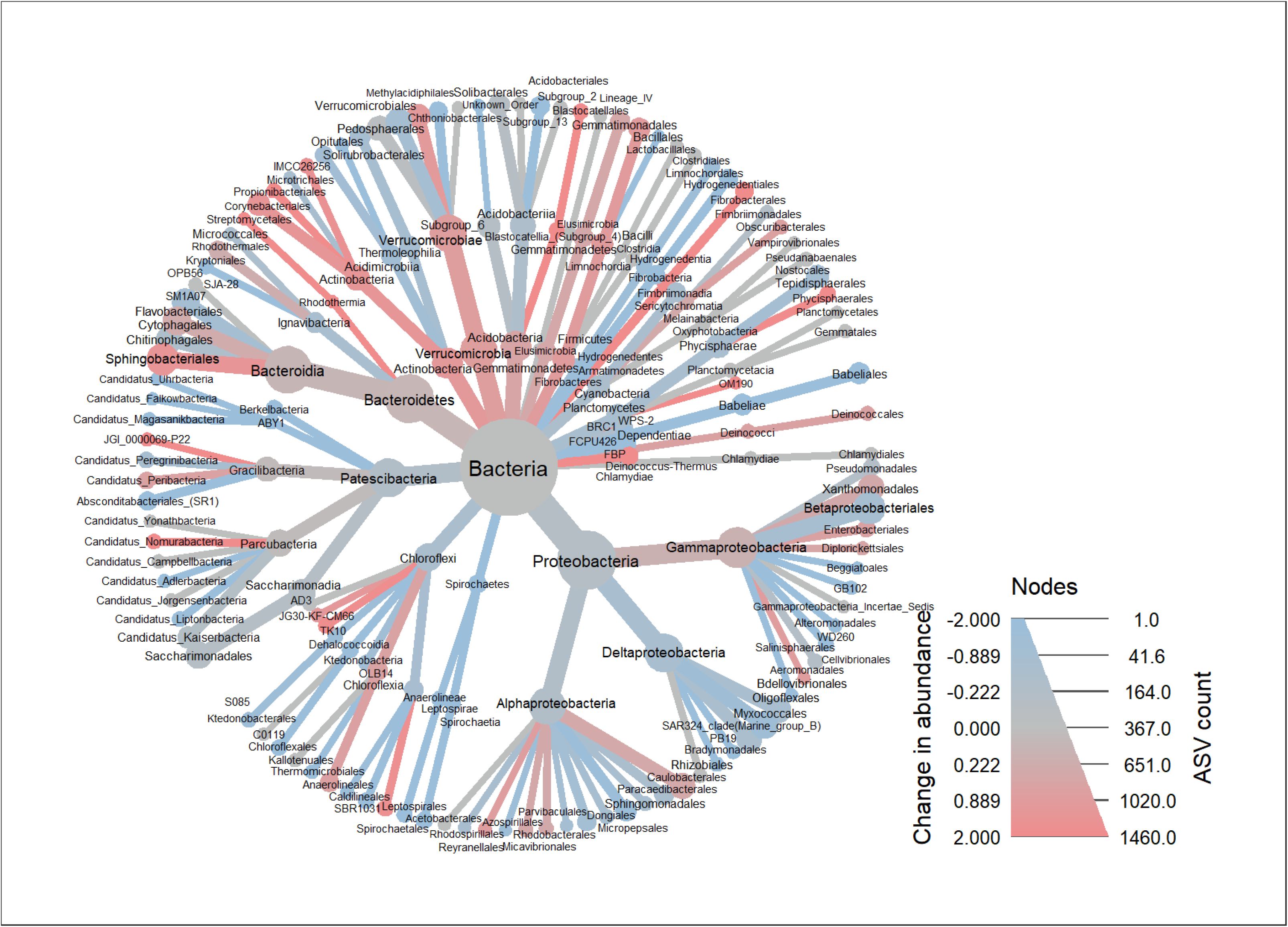
A heat tree representing the log_2_ differences in relative abundances of the rhizosphere bacteria from the B73 inbred line and the Ancho - More 10 Mexican landrace analysed in GH1. Taxa in higher abundances in B73 are in red, and in blue if in higher abundances in the Ancho - More 10 landrace.

Bulk soil was also taken before and after the 12-week sampling period and compared to the rhizosphere of the two landraces. Significant differences were found over the 12-week period in both community composition (**Fig S2A**) and richness (**Fig. S2B**). Post-hoc analysis of compositional differences revealed that soil did not vary from itself during the 12-week period (**Table S4**), and B73 line and Ancho - More 10 landrace did not vary from each other in this analysis. However both differed from the initial starting soil. B73 also significantly varied from the week-12 soil, while the Ancho - More 10 did not. Post-hoc analysis of richness revealed that all samples increased in richness from the initial soil samples (**Table S4**), but the week-12 soil and rhizosphere samples did not significantly differ from each other. Initially, there were 102.9 ASVs in soil at the start of the experiment, which rose to 236.3 ASVs after 12-weeks (that was non-significantly higher than in both varieties).

### GH2 – There were further significant differences between the rhizosphere bacteria of other landraces

Within GH2, the B73 inbred line and Ancho - More 10 landrace were supplemented with an additional four Mexican landraces (Apachito - Chih 172, Tehua - Chis 204, Serrano - Pueb 180 and Hairnoso de Ocho) and grown again over a 12-week period. As before, significant compositional differences were observed (*R*^2^ = 0.291, *P* = 0.001), with differences between varieties accounting for 29.1% of community variation (**Fig. 4A**). Since the B73 inbred line is so highly bred, B73 samples were removed to investigate for differences between Mexican landraces. Similarly, there were similarly significant differences in rhizosphere bacterial communities (*R*^2^ = 0.305, *P* = 0.001), accounting for 30.5% of community variation. Post-hoc analysis was subsequently performed to compare all 5 landraces, the B73 line and week-12 soil samples. There were significant differences between the Tehua - Chis 204 and C604 landraces specifically, but also between the week-12 soil communities and Tehua - Chis 204. While there was clear variation in the rhizosphere bacteria observable within families, there were also consistent differences between landraces (**Fig. 4C**). For example, within the B73 landrace, there were higher than average Devosiaceae, while in the Ancho - More 10 the Burkhoderiaceae were generally lower abundance than the other landraces.

**Figure 4.**
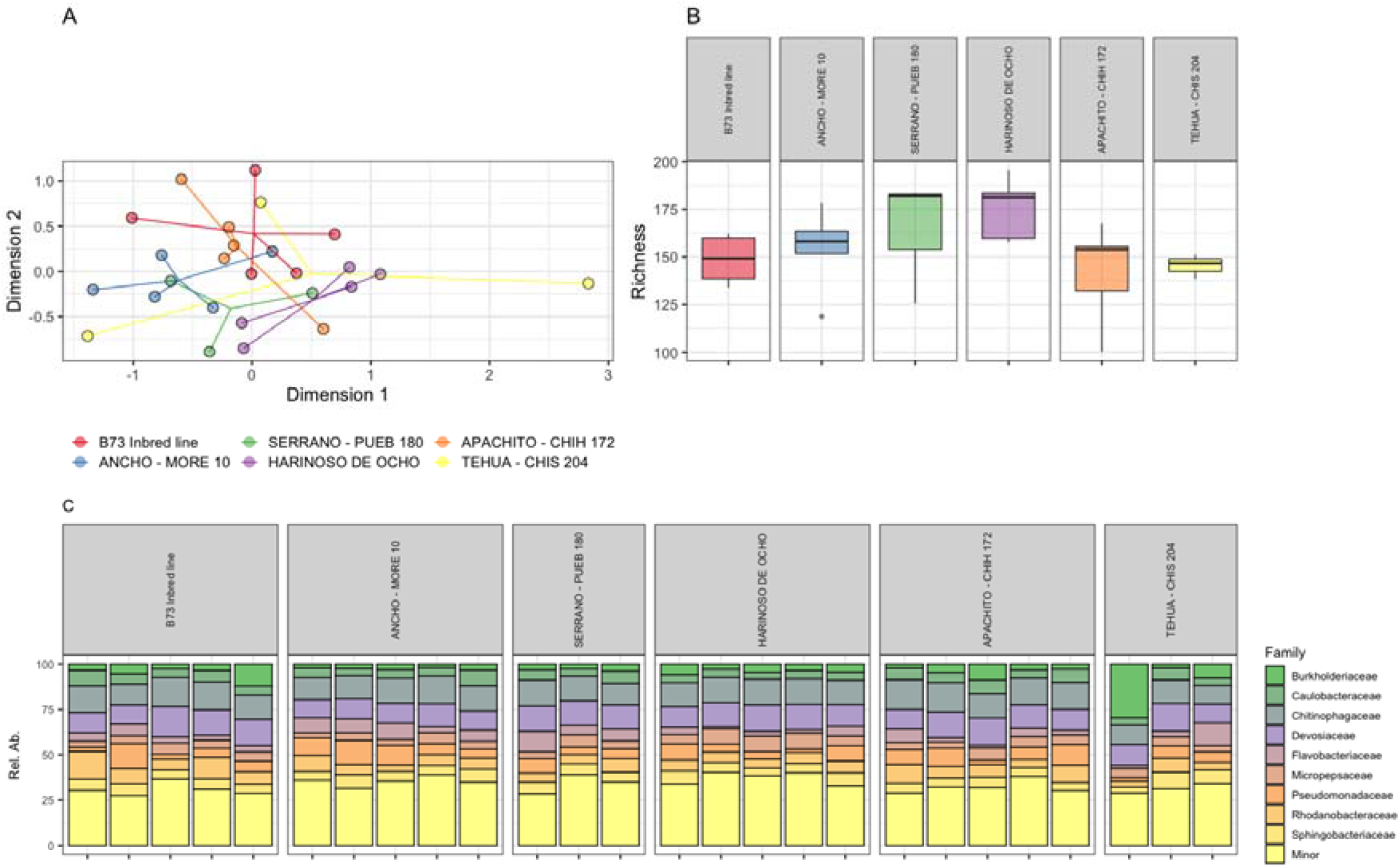
(**A**) Non-metric multidimensional scaling plot visualised significant differences between the rhizosphere bacteria from the six landraces from GH2. (**B**) While there were more ASVs on average within the C1580 and Hairnoso de Ocho landraces, they were non-significant. (**C**) The relative abundances of families significantly varied between landraces.

As before, there were visible differences in bacterial richness between the inbred line and landraces, with Ancho - More 10 (154.1 ± 22.0 ASVs) being higher than B73 (148.8 ± 12.5 ASVs) (**Fig. 4B**). The Hairnoso de Ocho landrace was the most bacterial rich (175.6 ± 16.3 ASVs), followed by Serrano - Pueb180 (163.8 ± 33.1 ASVs), Ancho - More 10, B73, and Apachito - Chih 172 (141.8 ± 26.5 ASVs), with Tehua - Chis 204 (129.3 ± 32.6 ASVs) being the least species rich. The genotypes screened in GH1 therefore had intermediate levels of bacterial richness compared to the other landraces. As before, these were not significantly different within a Kruskal-Wallis test (*H* = 2.08, *P* = 0.149) or post-hoc Dunn tests. Similarly, there were also no significant differences in ASV richness between landraces when the B73 inbred line was removed (*H* = 2.08, *P* = 0.149). Meanwhile, soil had the lowest starting ASV richness (97.4 ASVs) that rose during the 12-week incubation period (187.5 ± 4.9 ASVs), although as in GH1, this was also non-significantly higher (**Table S5**).

### GH2 – Bacterial community composition clustered with phylogenetic relatedness

A pairwise-distance matrix was estimated from the genomic data (DARTSEQ) of the genotypes produced by CIMMYT. Since the B73 inbred line has undergone intense breeding programmes, differences between it and the Mexican landraces may have driven the genome-microbiome correlation. Therefore the B73 was removed to leave only comparisons between Mexican landraces. A Bray-Curtis dissimilarity matrix produced from the GH2 rhizosphere bacteria was then correlated against this matrix using a Mantel statistic (*Z* = 35.4, *P* = 0.001) and Procrustes analysis (Sum of squares = 0.00, *P* = 0.001), both of which found a highly significant correlation between the two. A principal component analysis of the genomic matrix was performed and PC1 was plotted on the nMDS of GH2 to visualise this relationship (**Fig. S4**).

### Comparing the bacterial communities of GH1 and GH2

The B73 and Ancho - More 10 samples from GH2 were taken and compared to the data produced in GH1. There was no significant difference in the rhizosphere bacteria between genotypes when analysed together (*R*^2^ = 0.035, *P* = 0.636), however there was a large difference in between the bacterial communities from the two experiments (*R*^2^ = 0.594, *P* = 0.001), accounting for 59.4% of community variation (**Fig. S5**). The rhizosphere bacteria of GH1 had higher richness than GH2 on average, while Ancho - More 10 had higher richness than the rhizosphere of B73 with both experiments. However, neither genotype (*H* = 0.70, *P* = 0.402) nor experiment (*H* = 3.16, *P* = 0.076) had significantly different richness.

The bacteria of the bulk soil was also compared between the GH1 and GH2 samples to determine whether the inoculants from the surrounding soils differed. However, comparisons were limited due to a lack of replicates of the starting soil from GH2 as there were generally higher failure rates for soil samples than for the rhizosphere samples. However, there was evidence of a batch effect on the composition (**Fig. S7A**) and richness (**Fig. S7B**) of the bulk soil between experiments.

## Discussion

Results here confirm that there are differences in the rhizosphere bacteria between maize of different lineages. Furthermore, the rhizosphere bacterial community composition was correlated to the relatedness of the different varieties, finding that more closely related genotypes had more similar rhizosphere bacteria. For example, the Serrano – Pueb 180 and Ancho – More 10 landraces aggregated both in terms of its genomic distance and rhizosphere bacteria. This suggests that genomic variation between landraces influences the formation of the rhizosphere microbial assembly. It should be noted that this was performed with small sampling sizes, and interpretation of these results should therefore be taken with caution. Nonetheless, varieties were found to significantly differ in community composition in both the GH1 and GH2 experiments, while the Ancho – More 10 landrace also had higher richness than the B73 line in both GH1 and GH2 (albeit non-significantly so), and are worthy of further exploration. Other studies of the maize rhizosphere microbiome have been performed in experimental field plots (9–11,40) which have similarly found significant differences in the bacterial communities between differing landraces and inbred lines. Furthermore, a phylogenetic correlation between host and the rhizosphere bacteria was observed within the field between modern lines within the field (40). However, the only study to date performed on maize within glasshouses was performed using older technologies (DGGE and microarrays) that may have limited resolution (11). This work expanded on these previous studies in that it was performed in a controlled environment, and thereby limiting the potential of differences in micro-climate and edaphic properties, and it was also analysed using high-throughput sequencing techniques to profile both the bacterial community and the maize genomes in higher resolution. Together, these results confirm that differences between maize varieties are at least partially linked to genomic variation, and their effect size can be substantial enough to be observable within environmental settings.

Genotypic variation has been found to be relatively small compared to other factors such as environmental variation (10,17). In this work, we found the rhizosphere bacterial communities of the same varieties from GH1 and GH2 to be highly divergent, and to such an extent that the differences between the B73 and Ancho - More 10 communities were entirely confounded. While the starting soil was the same type, it was different batches used in GH1 and GH2, and this together with seasonal variation in temperature and exogenous light, and in the surrounding microbes within the glasshouse, resulted in a substantially different rhizosphere assembly. This further suggests that the differences in community variation associated with genotypic variation are small compared to environmental variation. There were also temporal differences in the bulk soil microbial communities from GH1 and GH2 (between the week-0 and week-12 samples). As in other studies (41,42), the rhizosphere also had lower bacterial richness than bulk soil after 12-weeks of incubation. We therefore further suggest that differences in rhizosphere bacteria between varieties are derived from differences in recruitment (either actively or passively) of local bacteria rather than the culturing of specific bacteria (43). This would also explain the lack of genomic variation found in other studies, and the considerable overlap between assemblages over environmental gradients (44). It also suggests that to fully understand the rhizosphere community assembly, the effect of the interaction between genome and environment needs to be considered.

The mechanisms that translate genomic variation between landraces into variation in the microbiome are also unknown. In this work, soil was equilibrated for, and seeds were likely to have nominal quantities of bacteria associated with them (since reads were dominated by host DNA). Therefore, differences in the initial bacterial inoculants were unlikely to have induced differences between landraces/lines within experiments. Differences in root architecture were observed between landraces from GH2 (experimental observations), and are one mechanism which might differentiate their associated microbial communities, but there is a plethora of other mechanisms that may have induced our observed differences. Differences in rhizoexudates have been shown to influence the rhizosphere microbiome of maize specifically (45), representing a mechanism worthy of further exploration. The Benzoxazinoids (BXs) are root exudates from maize that are particular interest since they have been shown to be notably higher in domesticated lines. They have been shown to offer protection against pathogenic microbes, insects and plants (45–48). Contrastingly, they have also been shown to recruit the plant growth-promoting bacterium *Pseudomonas putida* in the rhizosphere (45,49). The differential expression of BXs represents one pathway in which the rhizosphere bacteria are likely differentiated between landraces, and how domestication has shaped the rhizosphere microbiome. A comprehensive exploration of the genes associated with variation in the composition and rates of secretion of rhizoexudates would likely yield informative results. Ultimately, future studies would benefit from implementing a multi-omic approach to begin to unravel genotypic variation of the microbiome, with the integration of metagenomic, metatranscriptomic and metabolomic methods coupled with morphological observations.

While only small differences in the microbiome were observed between landraces in this study, the wider implications for host and ecosystem functioning remain unknown. The colonisation of single pathogens can induce diseases that severely reduce host vitality (50). Conversely, the composition and abundances of arbuscular mycorrhizal fungi have been shown to have differing growth benefits within crops *in situ* (51). It is therefore feasible that the small differences in commensal microbes observed within this study have biologically meaningful effects on the host functioning and are worthy of further exploration. Results here suggest that domestication has almost certainly influenced the composition of the maize rhizosphere microbiome, which may or may not have influenced its functioning. Throughout the domestication process, traits were selected for including yield and drought resilience (42). However, it is possible that a proportion of these traits arose through the selection of the microbiome. The study of the genome-microbiome interaction through the domestication process of maize would therefore be of particular interest.

