## Supplementary material for "THE RHIZOSPHERE BACTERIAL COMMUNITIES DIFFER AMONG DOMESTICATED MAIZE LANDRACES – AN EXPERIMENTAL CONFIRMATION": Figure S1-S7

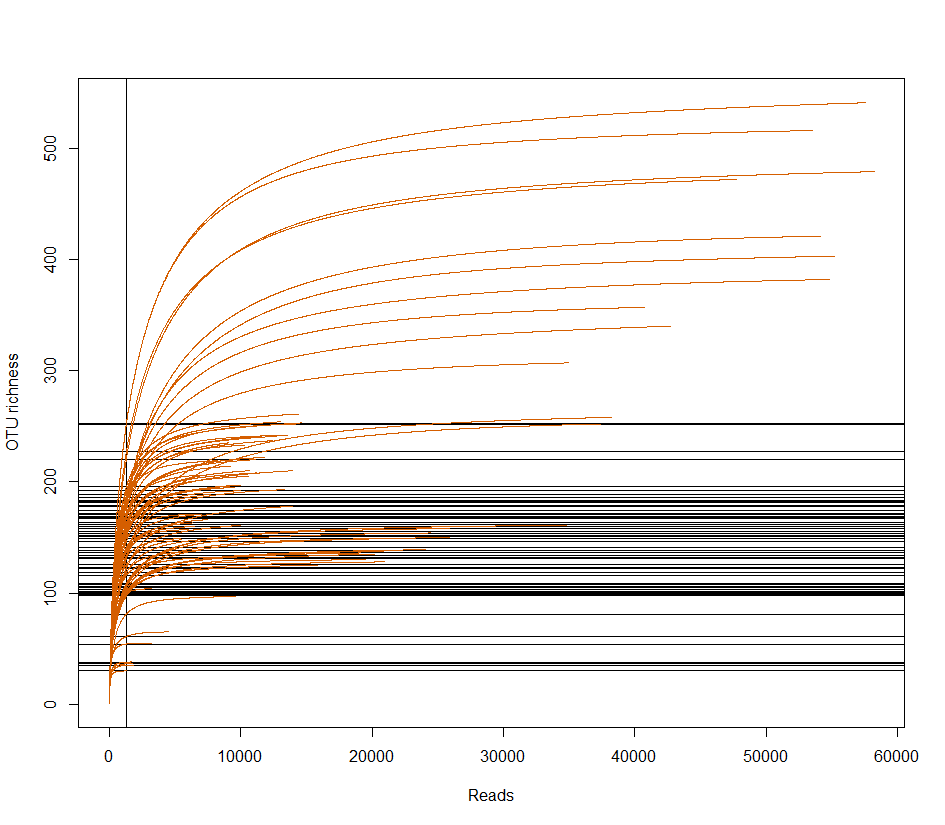

**Figure S1** Rarefaction curves suggest that an asymptote were almost reached for all samples. The minimum read depth was 1307 reads.

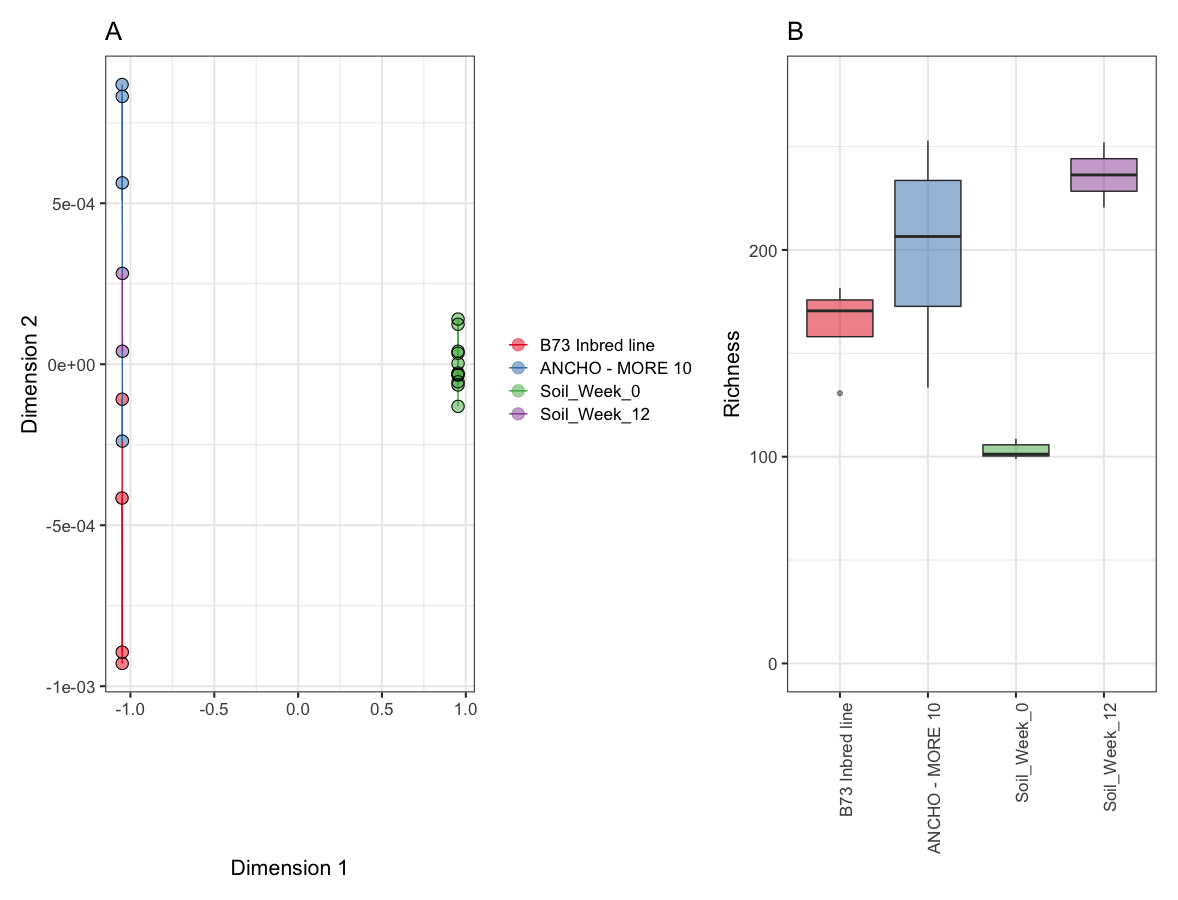
**Figure S2** (**A**) Non-metric multidimensional scaling plot visualised significant differences between the rhizosphere bacteria from the B73 and C233 landraces from GH1. (**B**) While there were more ASVs on average within the C233 landrace, it was non-significant.

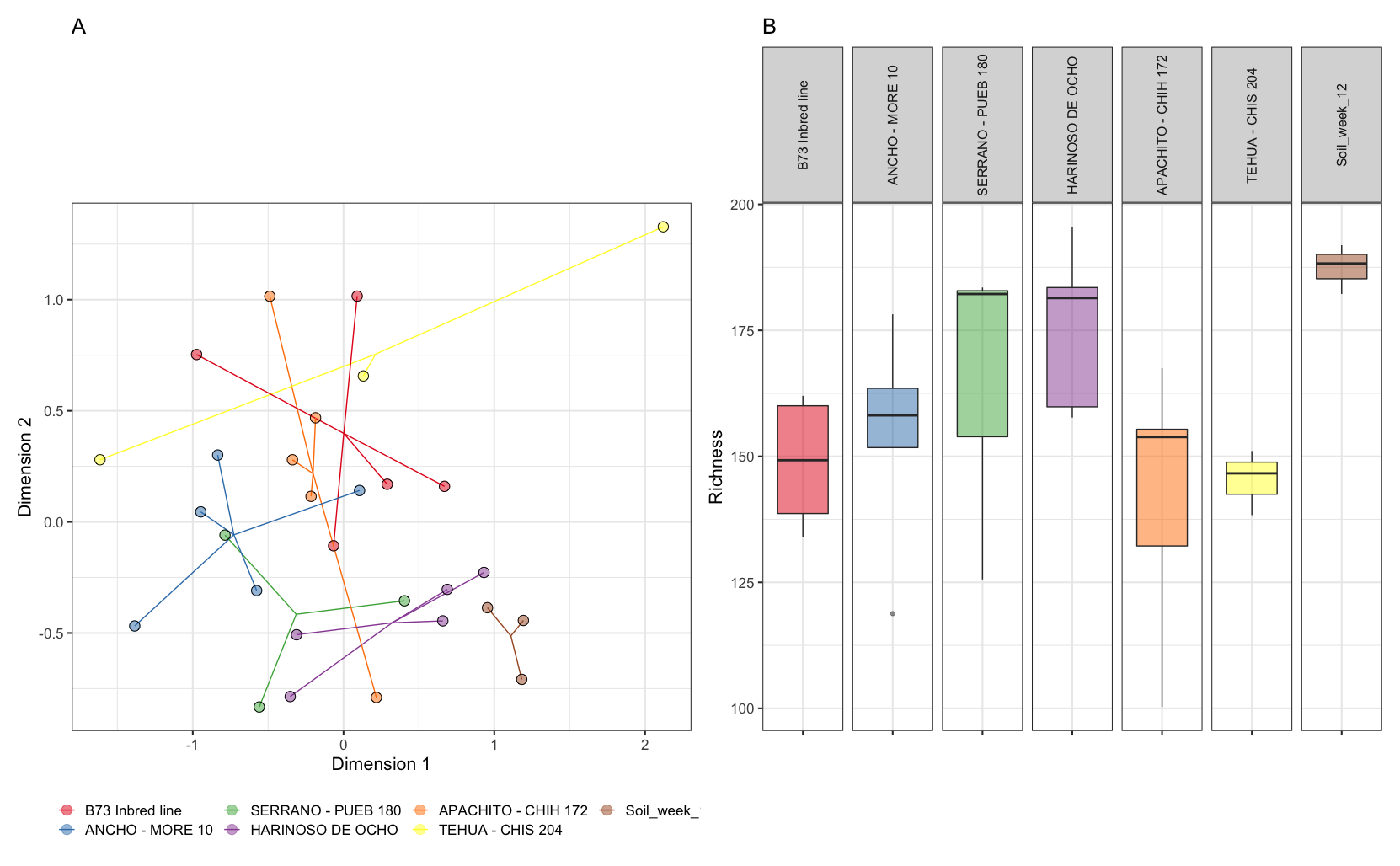

**Figure S3** (**A**) Non-metric multidimensional scaling plot visualised significant differences between the rhizosphere bacteria from the six landraces and bulk soil from GH2. (**B**) While there were more ASVs on average within the SERRANO – PUEB 10 and HARINOSO DE OCHO landraces, they were non-significant. Due to multiple failures, soil-0 were excluded from plots.

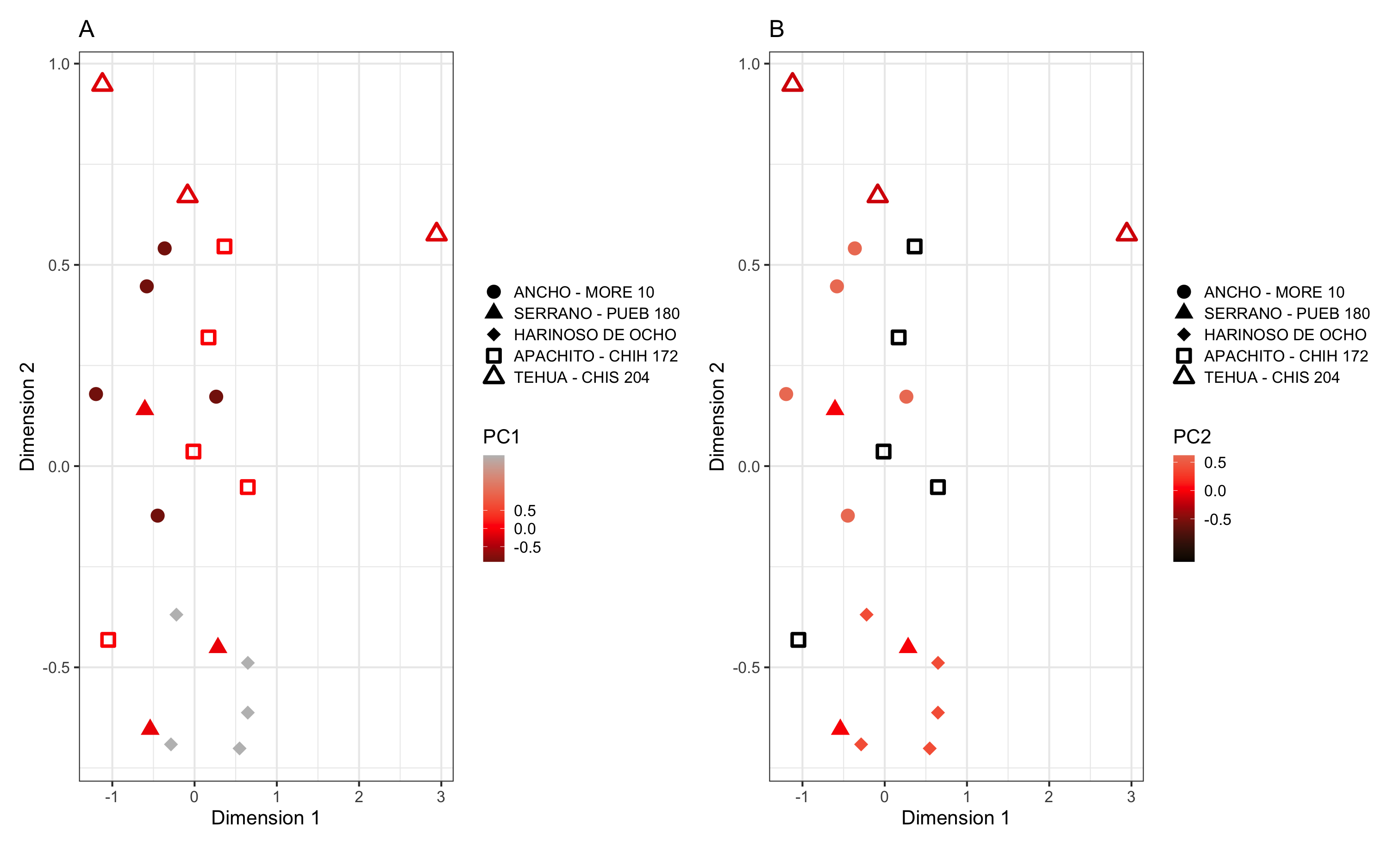
**Figure S4** Principal component analysis was performed on the genomic matrix of the different landraces. The first principal component (PC1) (A) and second principal component (PC2) (B) were subsequently plotted on the nMDS plot of the rhizosphere bacteria from GH2.

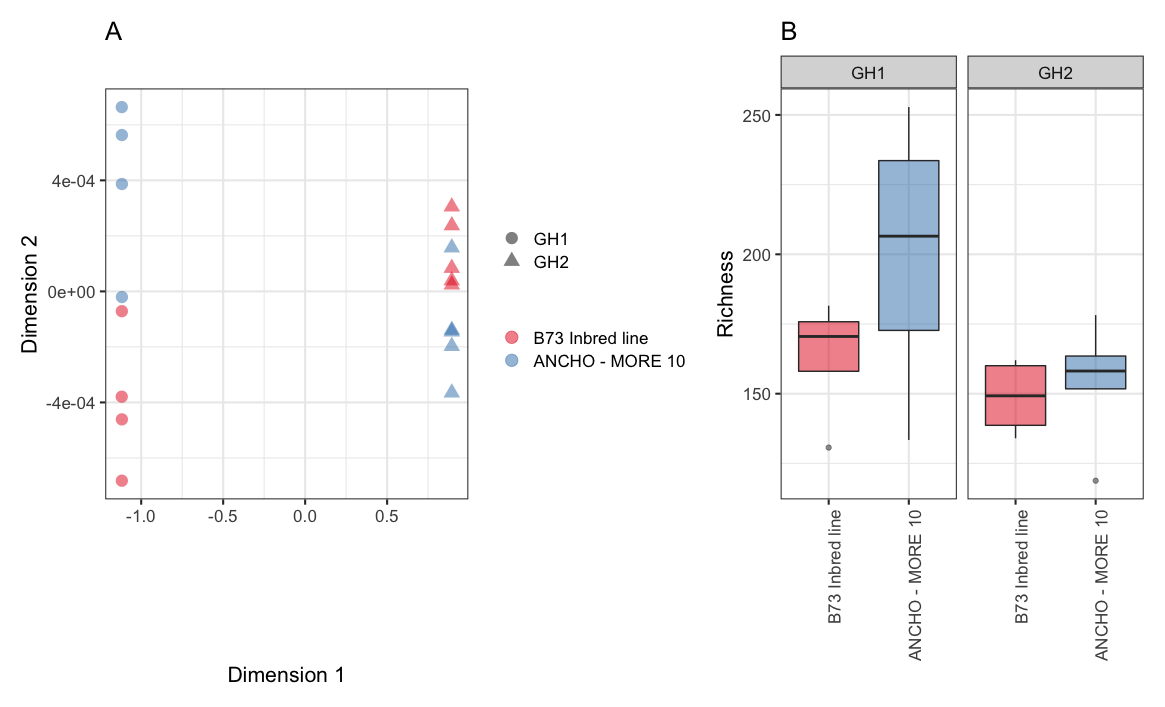
**Figure S5** The B73 and ANCHO - MORE 10 samples from GH1 and GH2 were analysed together. (**A**) Non-metric multidimensional scaling plot demonstrated a large effect of the different experimental setups. (**B**) Similarly, while the ANCHO – MORE 10 landrace was higher than the B73 landrace, the samples from GH1 were higher in bacterial richness than the samples from GH2.

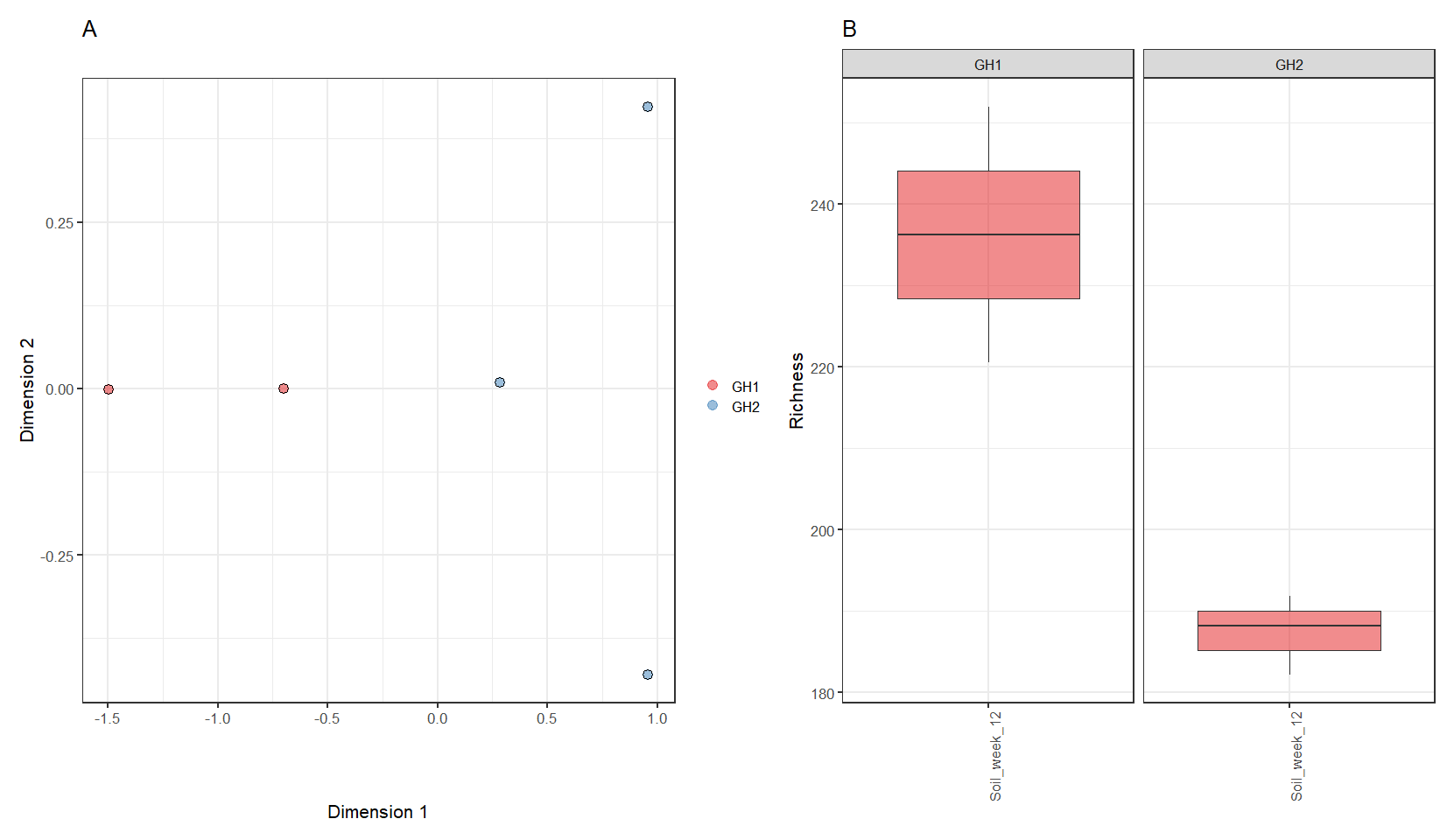
**Figure S6** Bulk soil samples from GH1 and GH2 were compared. Due to low sampling replicates, only samples from after the 12-week incubation period were compared. (**A**) Non-metric multidimensional scaling plot demonstrated a large effect of the different experimental setups. (**B**) Similarly, there was a notable difference in ASV richness of the bulk soils from the GH1 and GH2 samples taken after 12-weeks of incubation.
